# Cognitive process modeling addresses context independence violations in the ABCD Study stop-signal task

**DOI:** 10.1101/2021.07.26.453872

**Authors:** Alexander Weigard, Dora Matzke, Charlotte Tanis, Andrew Heathcote

**Author notes:** Correspondence concerning this article should be addressed to Alexander Weigard, University of Michigan Department of Psychiatry, Rachel Upjohn Building, 4250 Plymouth Road, Ann Arbor, MI 48109.

## Abstract

The Adolescent Brain Cognitive Development (ABCD) Study is a longitudinal neuroimaging study of unprecedented scale that is in the process of following over 11,000 youth from middle childhood though age 20. However, a design feature of the study’s stop-signal task violates “context independence”, an assumption critical to current non-parametric methods for estimating stop-signal reaction time (SSRT), a key measure of inhibitory ability in the study. This has led some experts to call for the task to be changed and for previously collected data to be used with caution. We present a formal cognitive process model, the BEESTS-ABCD model, that provides a mechanistic explanation for the impact of this design feature, describes key behavioral trends in the ABCD data, and allows biases in SSRT estimates resulting from context independence violations to be quantified. We use the model to demonstrate that, although non-parametric SSRT estimates generally preserve the rank ordering of participants’ SSRT values, failing to account for context independence violations can lead to erroneous inferences in several realistic scenarios. Nonetheless, as the BEESTS-ABCD model can be used to accurately recover estimates of SSRT and other mechanistic parameters of interest from ABCD data, the impact of such violations can be effectively mitigated.

## Introduction

Response inhibition, the ability to stop prepotent responses or actions that are no longer contextually appropriate, is thought to be a foundational component of self-control (Miyake et al., 2000; Ridderinkhof et al., 2004; Verbruggen et al., 2014) and has long been of key interest in research on multiple clinical conditions, including Attention-Deficit/Hyperactivity Disorder (ADHD) and problematic substance use (Boonstra et al., 2010; Gorenstein & Newman, 1980; Mahmood et al., 2013; Nigg, 2017; J. L. Smith et al., 2014). The stop-signal paradigm (Logan et al., 1984; Verbruggen & Logan, 2008), one of the most widely employed laboratory measures of response inhibition, probes this ability in the context of a two-choice decision task. On a minority of “stop” trials a visual or auditory “stop signal”, which indicates that participants must withhold their response on that trial, is presented after a variable delay following the onset of the choice stimulus (the “stop-signal delay” or SSD). Depending on both the SSD and the participant’s inhibitory ability, either inhibition fails and a choice response is made (a “signal response”) or the response is successfully withheld.

An appealing feature of the stop-signal paradigm is that it was explicitly designed with a cognitive model in mind that can be leveraged to precisely measure inhibitory ability: the “independent race model” (Logan, 1994; Logan et al., 1984; Logan & Cowan, 1984). This model posits that, on stop trials, a “go” process triggered by the choice stimulus races a “stop” process triggered by the stop signal. When the stop process wins the race, the response is inhibited, whereas inhibition fails when the go process finishes first. By making only limited assumptions about the distributions of times to complete these processes (e.g., symmetry, or that the stop process takes a fixed time) and the effect of the stop signal (e.g., “context independence”; that the go process is the same regardless of whether or not there is a stop signal), the speed of the latent stop process, or “stop-signal reaction time” (SSRT), can be estimated (Logan, 1994; Matzke et al., 2018). Such “non-parametric” SSRT estimates are used to measure individuals’ response inhibition ability across many neuroscientific and clinical research applications [e.g., (Aron & Poldrack, 2006; Lipszyc & Schachar, 2010; Nigg et al., 2006)].

Given this task’s popularity and the ability of SSRT to precisely index the integrity of response inhibition, it is not surprising that the task has been included in one of the most ambitious research efforts of our time: the Adolescent Brain Cognitive Development (ABCD) Study (Casey et al., 2018; Garavan et al., 2018). The ABCD study is a multi-site collaboration in the United States that has recruited a diverse a sample of over 11,000 9- and 10-year-old children and aims to follow them prospectively through at least age 20 to acquire a rich array of longitudinal data from neuroimaging, cognitive, personality, psychiatric and sociocultural domains. Although initially conceived with the goal of assessing the impact of substance use on adolescents’ brains (Volkow et al., 2018), ABCD has grown into an unprecedented interdisciplinary collaboration and open data source that is beginning to drive new insights in areas as diverse as network neuroscience (Marek et al., 2019; Sripada et al., 2019), child psychopathology (Clark et al., 2021; Funkhouser et al., 2020; Mennies et al., 2020), and bilingualism (Dick et al., 2019). As several lines of work now underscore the importance of large, population-based samples for bolstering the reproducibility of behavioral and neuroscientific research (Button et al., 2013; Etz & Vandekerckhove, 2016; Falk et al., 2013; Loken & Gelman, 2017; Marek et al., 2020; Poldrack & Gorgolewski, 2014; Szucs & Ioannidis, 2020), the ABCD Study presents a critical opportunity for scientists to characterize the clinical and neural correlates of response inhibition and related neurocognitive functions.

However, concerns have recently been raised that the design of the ABCD stop-signal task invalidates non-parametric SSRT estimates (Bissett et al., 2021). In the ABCD task, which is completed during functional magnetic resonance imaging (fMRI), children are presented right- or left-facing arrows and instructed to respond with the direction of the arrows via a button box. On stop trials (60 of 360 total trials), an upward-facing arrow, serving as a visual stop signal, replaces the choice stimulus at an SSD determined by a staircase algorithm: increasing or decreasing by 50ms depending, respectively, on whether or not the last inhibition was successful (Logan, 1994). Replacement limits the availability of the choice stimulus to the SSD duration. For example, when SSD = 0, the choice stimulus is never presented, so signal-respond choice accuracy is necessarily at chance. Choice accuracy increases with SSD to an asymptotic level at SSDs of approximately 0.3-0.4s (Bissett et al., 2021). In contrast, on “go” trials (choices without a stop signal) the presentation time of the choice stimulus is set to the shorter of 1s or the participant’s response, so the information necessary for accurate responding is equally present on every go trial. This constitutes a violation of “context independence”, as the go process is clearly not the same across go and stop trials. Crucially, violations of this assumption can bias non-parametric estimates of SSRT, leading Bissett et al. (2021) to suggest that the ABCD stop-signal task be changed and that previously collected data be interpreted with caution.

Although this design feature presents specific problems for SSRT estimation in the ABCD Study, it is important to note that the non-parametric approach is also unable to address a more general problem that potentially affects all stop-signal tasks: “trigger failure”, in which inattention prevents the stop signal from triggering the inhibitory process, precluding it from entering the race. Trigger failures have long been recognized as a possibility (Band et al., 2003; Logan, 1994), but their prevalence has only recently been determined using new, parametric approaches. Such approaches use cognitive-process models to provide detailed descriptions of the latent processes that contribute to task performance and allow measurement of these processes through parameter estimation. Matzke and colleagues (Matzke, Love, et al., 2017) extended the Bayesian parametric estimation of stop-signal reaction time distributions (BEESTS) framework (Matzke et al., 2013) to estimate the probability of trigger failures. They showed that failing to account for trigger failures can significantly bias non-parametric SSRT. Subsequent work found that trigger failures largely accounted for poor stop-signal task performance in schizophrenia (Matzke, Hughes, et al., 2017) and ADHD (Weigard et al., 2019) that had previously been attributed to inhibitory deficits due to the bias in non-parametric SSRT estimates. Hence, trigger failure illustrates the key advantages of a parametric approach, grounded in cognitive-process modeling, for describing the array of factors that contribute to stop-signal task performance and allowing their distinct contributions to effects of interest to be precisely measured.

Here we combine BEESTS with evidence accumulation modeling (Donkin & Brown, 2018), one of the most widely applied and validated cognitive model classes, and leverage previous models of visual-masking effects on perceptual choice (Ratcliff & Rouder, 2000; P. L. Smith & Ratcliff, 2009; P. L. Smith & Sewell, 2013) to describe how replacement of the choice stimulus by the stop signal impacts performance on the ABCD task. We show that this new cognitive-process model (BEESTS-ABCD) provides an accurate description of key trends in ABCD stop trial data, including the SSD effect on signal-respond choice accuracy that is the hallmark of the context independence violation. We next use BEESTS-ABCD to demonstrate that this violation can, if unaccounted for, lead to consequential biases in non-parametric SSRT estimates. Crucially, we show that our model can overcome these measurement difficulties, allowing existing data from the ABCD sample to be used to obtain unbiased estimates of SSRT (and other processes of interest) and allowing the same task design to continue to be used in future ABCD data collection, maintaining longitudinal comparability.

## The BEESTS-ABCD Model

BEESTS models the times for each of the go and stop process runners to complete its race as an ex-Gaussian distribution, a form that provides an excellent descriptive account of response time (RT) distributions (Heathcote et al., 1991). It is the sum of a normal distribution (with mean, *μ*, and standard deviation, *σ*) and an exponential distribution (with mean *τ*), where the latter produces the positive skew characteristic of RT distributions. Parametric models such as BEESTS do not need to assume context independence in order to provide valid SSRT estimates when they are fit to only stop trial data (Matzke et al., 2021), but usually do so in order to leverage information provided by go trials to make estimation more efficient. Although estimating parameters from go and stop trials separately could, in principle, address context independence violations in ABCD, such estimates are unlikely to be sufficiently reliable because the design has relatively few stop trials. Fortunately, evidence accumulation processes can be incorporated into the BEESTS framework to provide a unified account of the ABCD choice task on go and stop trials, enabling reliable and valid estimation of both SSRT and trigger failures.

There are a variety of different evidence accumulation models, including some compatible with a race-model framework [e.g., (Brown & Heathcote, 2008; Tillman et al., 2020)], but all assume choices are made by gradually collecting evidence for each option until a threshold (*B*) amount is obtained for a given option, triggering the corresponding response. RT is the sum of the time to reach threshold and non-decision time (*t_0_*), which is made up of the time to initially encode the stimulus into a form suitable for obtaining decision-relevant evidence and the time to produce a motor response. For briefly presented visual stimuli that are followed by a mask, the encoding process is assumed to establish a representation in visual short-term memory that persists after the choice stimulus disappears. The strength of this representation, which determines the rate at which evidence accumulates (*v*), increases with presentation duration up to some asymptotic level (Ratcliff & Rouder, 2000; P. L. Smith & Ratcliff, 2009; P. L. Smith & Sewell, 2013).

Evidence can be modeled as the sum of two parts, a *discriminative* component, which differentially favors the option that matches the stimulus over the option that mismatches the stimulus, and an *urgency* component, which favors each option equally and ensures that even very difficult choices produce a timely response (Mazurek et al., 2003; van Ravenzwaaij et al., 2019). We assume that only urgency (with associated rate *v*_0_) is present when SSD = 0 and that the discriminative component gradually grows as SSD increases and the visual short-term memory representation strengthens. By assuming that asymptotic discrimination (likely achieved around 0.3-0.4s SSDs where choice accuracy is also asymptotic) and urgency are the same for go and stop trials, the model achieves efficient estimation by leveraging information obtained on the more common go trials to constrain estimates for stop trials.

BEESTS-ABCD’s evidence accumulation component has the same independent-race architecture as BEESTS (Matzke et al., 2019), with one accumulator (“racer”) corresponding to each potential response. We estimate rates *v*_+_ and *v*_-_ for accumulators that, respectively, match and mismatch the presented stimulus. Following a previously proposed stop-signal model (Logan et al., 2014), we assume the accumulators are single threshold diffusion processes, where an additive infinitesimal Gaussian noise causes the evidence total to fluctuate during accumulation (Tillman et al., 2020). In contrast to Logan et al. (2014), we do not make the same assumption for the stop racer, as the resulting model is poorly identified and hence cannot be used for measurement purposes (Matzke et al., 2020).

Instead, we formed a hybrid of the model proposed by Logan et al. (2014) and the Matzke et al. (2019) BEESTS model by characterizing the finishing time of the stop racer with an ex-Gaussian distribution (Figure 1A). This avoids the cause of the estimation problems in Logan et al. (2014): having to estimate a non-decision time parameter for the stop racer that determines the finishing-time distribution’s lower bound^1^. This “hybrid” model can support accurate parameter estimation with practically achievable numbers of trials even in quite complex factorial designs (Tanis et al., in preparation). As illustrated in Figure 1B, we modeled the increase in discriminative information with longer presentation durations as linear increases and decreases, respectively, for matching and mismatching accumulators with the same absolute slope *g*, taking them both from *v*_0_ at SSD = 0 to their respective asymptotic values of *v*_+_ and *v*_-_ at longer SSDs.

**Figure 1.**
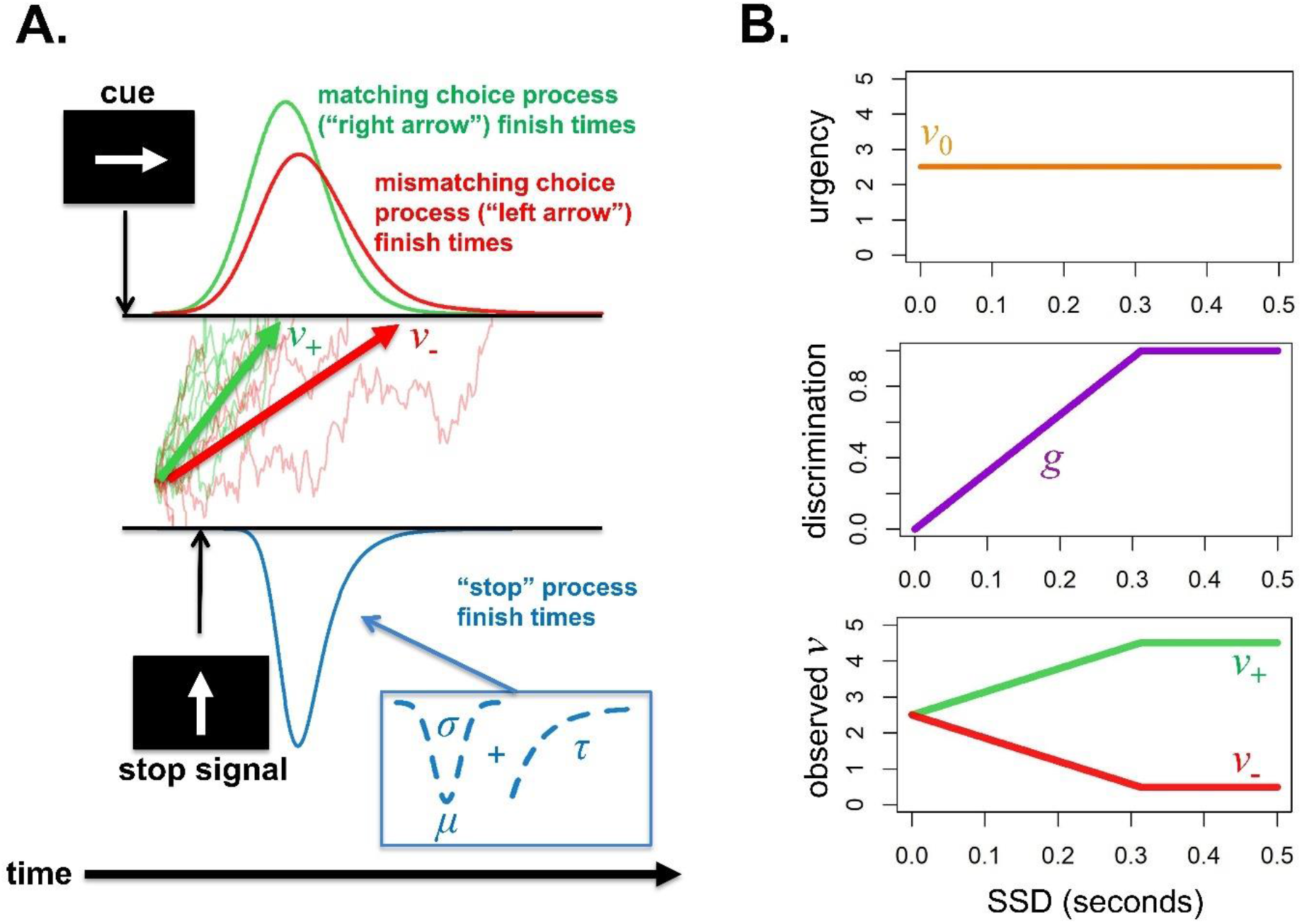
Key components of the BEESTS-ABCD model. (**A**) The “hybrid” modeling framework (Tanis et al., in preparation) that combines an evidence accumulation model of the “go” process with an ex-Gaussian model of “stop” process response times (RTs). Go process RTs result from a race between accumulators that gather noisy evidence for the choices matching and mismatching the stimulus (in this example, a right-facing arrow) at average rates of *v*_+_ and *v*_-_, respectively, until one accumulator crosses an upper response threshold. Stop process RTs are drawn from a Gaussian distribution specified by mean (*μ*) and standard deviation (*σ*) parameters and convolved with an exponential distribution with mean *τ*. (**B**) The model’s explanation for the impact of context independence violations on stop trials. Evidence signals for the matching and mismatching accumulators on stop trials of a given SSD are the sum of an *urgency* component that drives evidence accumulation for both choices equally and a *discrimination* component that favors the choice matching the presented stimulus. Urgency is determined by parameter *v*_0_ and is identical across all SSDs. The discrimination component is completely absent (equal to 0) at a 0s SSD, as the choice stimulus is not presented, but increases linearly at the same rate *g* for both matching and mismatching components until they reach the level of go trials. Therefore, the match and mismatch rates are identical on 0s SSD trials and gradually move apart from each other at longer SSDs until they become equal to their go trial levels.

In summary, BEESTS-ABCD has 11 parameters. Two accommodate the context-independence violation: urgency (*v*_0_) and the rate of growth in discriminative perceptual information (*g*). The remainder are shared with the original hybrid model (Tanis et al., in preparation): asymptotic matching (*v*_+_) and mismatching (*v*_-_) rates, non-decision time (*t*_0_) and threshold (*B*) for the choice accumulators, the ex-Gaussian parameters for the stop runner (*μ*, *σ*, *τ*), the probability of trigger failure (*p*_tf_) and the probability of failing to respond to the go stimulus (“go failures”, *p*_gf_). Although relatively rare, go failures were included as, like trigger failures, they have been shown to bias estimates of inhibition if neglected (Matzke et al., 2019; Tannock et al., 1995).

## Results

In this section, we: 1) show that BEESTS-ABCD displays excellent fit to stop-signal task data from the ABCD Study; 2) provide examples illustrating how unaccounted for variation in the context-independence-violation parameters can confound inferences about inhibitory ability; and 3) demonstrate that BEESTS-ABCD can be effectively used to estimate SSRT and other parameters of interest with existing ABCD data.

### Application to ABCD Stop-Signal Task Data and Model Fit

We fit the BEESTS-ABCD model to data from a randomly selected subsample (*n*=600) of ABCD participants using Bayesian estimation. A subsample, rather than the full ABCD sample, was analyzed because Bayesian estimation is computationally expensive. Priors for parameter estimation were informed by posteriors from a Bayesian hierarchical model fit to an independent subset of 300 individuals from the same ABCD Study sites. Procedures for creation of the subsamples, performance exclusion criteria, and model estimation are detailed in Materials and Methods.

Table 1 displays group average parameter estimates for the BEESTS-ABCD model. The 6 go parameters indicate that go failures are rare (2.7%), and that the mean times for matching and mismatching go accumulators to finish (*t_0_* + *B*/*v_+_* and *t_0_* + *B*/*v_-_*, respectively) are 0.57s and 3.67s. Correct responses associated with the matching accumulator winning have a mean RT of 0.55s, whereas error responses have a mean of 0.58s. In comparison, when there is no discriminative information (*v_0_* only), the mean finishing time for the go race is slightly faster, at 0.52s. The 3.7% trigger failure rate is on the low end of previous findings with non-clinical groups, indicating that participants were generally attentive to the task. The average SSRT estimated by the model is 0.268s, considerably slower than most previous BEESTS estimates for relatively easy go tasks in older participants (Matzke, Hughes, et al., 2017; Skippen et al., 2019), but comparable to estimates from children of similar age (0.243s for healthy 8-12 year-olds: Weigard et al., 2019).

**Table 1.**
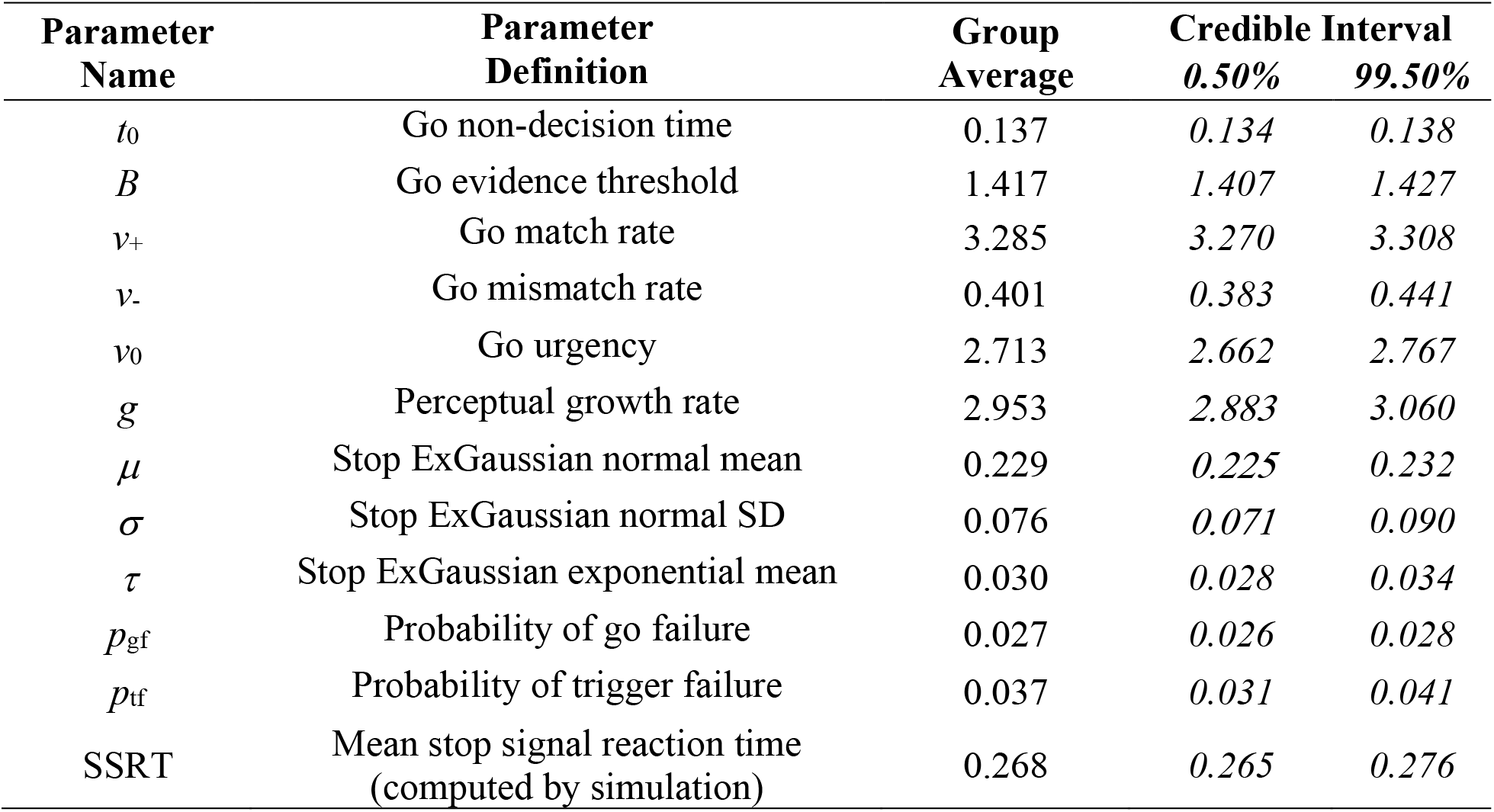
Average BEESTS-ABCD model parameter estimates and model-based mean stop signal reaction time (SSRT) estimates for the 600-participant group and the 99% posterior credible intervals for these group averages. The tight credible intervals for all parameters reflect the large (600 person) sample size on which the group average is based.

| Parameter Name | Parameter Definition | Group Average | Credible Interval |  |
| --- | --- | --- | --- | --- |
|  |  |  | 0.50% | 99.50% |
| $t_0$ | Go non-decision time | 0.137 | 0.134 | 0.138 |
| $B$ | Go evidence threshold | 1.417 | 1.407 | 1.427 |
| $v_+$ | Go match rate | 3.285 | 3.270 | 3.308 |
| $v_-$ | Go mismatch rate | 0.401 | 0.383 | 0.441 |
| $v_0$ | Go urgency | 2.713 | 2.662 | 2.767 |
| $g$ | Perceptual growth rate | 2.953 | 2.883 | 3.060 |
| $\mu$ | Stop ExGaussian normal mean | 0.229 | 0.225 | 0.232 |
| $\sigma$ | Stop ExGaussian normal SD | 0.076 | 0.071 | 0.090 |
| $\tau$ | Stop ExGaussian exponential mean | 0.030 | 0.028 | 0.034 |
| $p_{gf}$ | Probability of go failure | 0.027 | 0.026 | 0.028 |
| $p_{tf}$ | Probability of trigger failure | 0.037 | 0.031 | 0.041 |
| SSRT | Mean stop signal reaction time<br>(computed by simulation) | 0.268 | 0.265 | 0.276 |

Figure 2A shows how SSD in the ABCD stop trial design affects choice processing on average, and for 20 randomly selected individuals. Matching (*v*_+_) and mismatching (*v*_-_) accumulator rates grow from being identical at 0s SSDs to being equivalent to their go trial levels at SSDs of between 0.3s and 0.4s, consistent with the SSD region where accuracy becomes asymptotic (Bissett et al., 2021). Posterior predictive plots (Gelman et al., 1996), which represent model fit by comparing a Bayesian model’s predictions with key trends in empirical data, indicate that the BEESTS-ABCD model provides an excellent description of the increase in empirical choice accuracy with SSD (Figure 2B). In contrast, a “go independence” model – identical to the BEESTS-ABCD model except that the go process is context independent (i.e., rates do not change with SSD) – displays increasingly gross misfits to empirical accuracy as SSD decreases from .35s (Figure 2C). Therefore, BEESTS-ABCD’s perceptual growth process generates a pattern of context independence violations that can explain the hallmark effect of SSD on choice accuracy in ABCD data.

**Figure 2.**
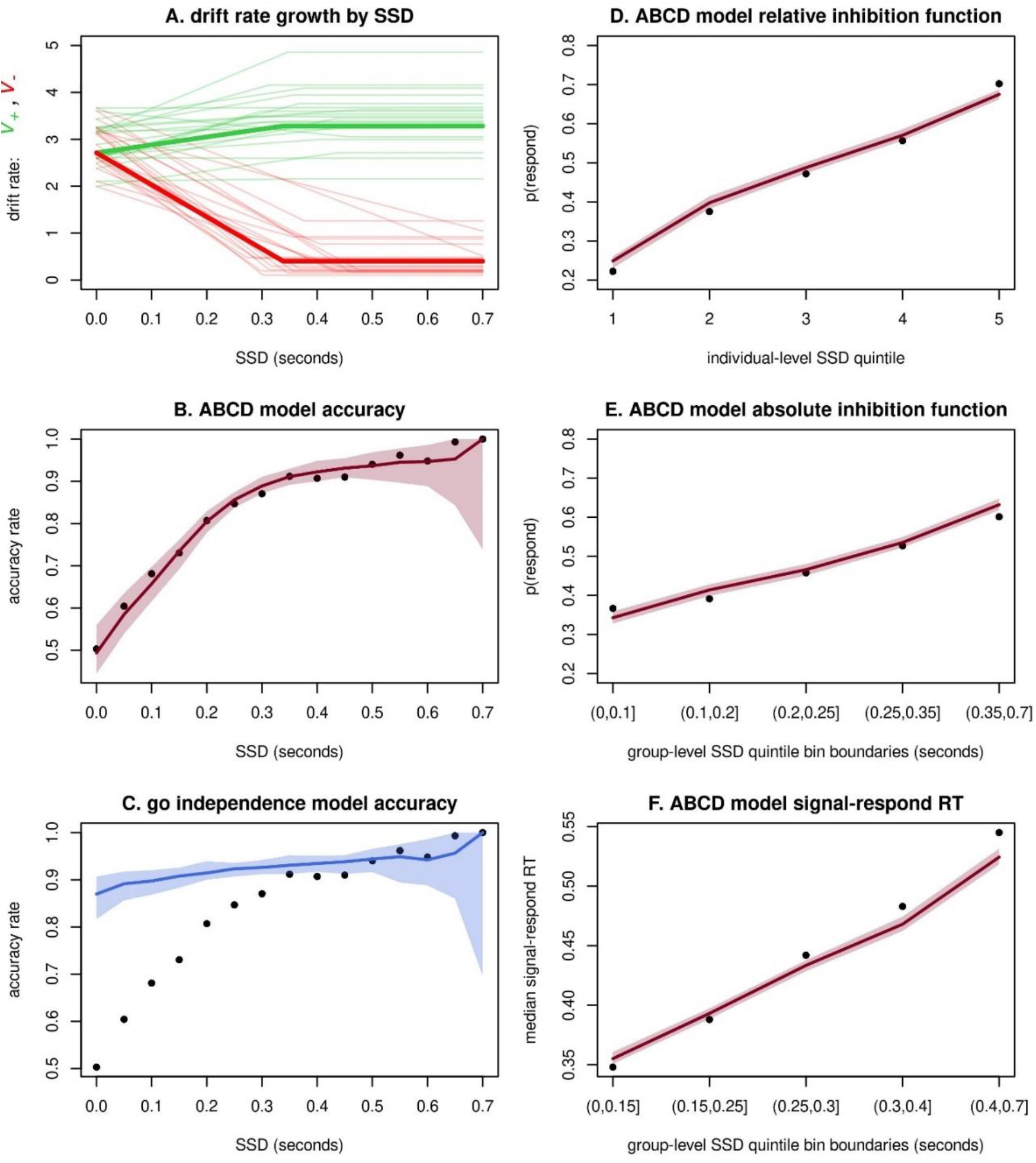
Results of the BEESTS-ABCD model application to a subsample of 600 ABCD participants. (**A**) Empirical growth patterns of matching (green lines increasing from SSD = 0) and mismatching (red lines decreasing from SSD = 0) “go” process accumulator rates by stop-signal delay (SSD) for the sample average parameter estimates (thick lines) and for parameter estimates from 20 randomly drawn participants (thin, faint lines) to illustrate individual variability. (**B**) The model’s median predictions (red line) and 99% credible interval (CI) of predictions (red shading) for accuracy rates across SSDs, overlayed with empirical values (dots). (**C**) Median (blue line) and 99% CI (blue shading) of the corresponding accuracy predictions of a “go independence” model, which assumes that the go process across all SSDs is identical to that on go trials, overlayed with empirical values (dots). (**D**) Median (red line) and 99% CI (red shading) of the model’s predictions for the “relative” inhibition function, for which SSDs are binned at the individual level to account for variation in performance, overlayed with empirical values (dots). (**E**) The model’s predictions for the “absolute” inhibition function, in which SSDs are binned across the group, overlayed with empirical values. (**F**) The model’s predictions for changes in median signal-respond response time (RT) by SSD with SSDs binned across the group, overlayed with empirical values.

We also sought to test whether BEESTS-ABCD explains other empirical trends that are conventionally used as benchmarks of goodness-of-fit in the stop-signal modeling literature (Matzke et al., 2018). The first is the pattern – predicted by the race model – in which the probability of responding increases with SSD. Following prior work (e.g., Weigard et al., 2019), we plotted this “inhibition function” using relative SSD bins based on quintiles (i.e., five equal groups of ordered SSDs) for each individual participant to account for individual variation in inhibitory performance (Figure 2D). Forming SSD bins based on absolute times confounds within- and between-participant performance (as different individuals have different ranges of SSDs due to the adaptive tracking algorithm), leading to a flatter inhibition function (Figure 2E). Regardless, in both cases the ABCD model provides a good description^2^.

The second trend is that signal-respond RTs tend to increase with SSD because the “go” RT distribution is censored by successful stopping to a lesser degree at longer SSDs (Colonius et al., 2001; Matzke et al., 2018). Following prior work (Weigard et al., 2019), we plotted this trend using “absolute” quintile bins calculated from SSDs collapsed across the signal-respond trials of all individuals. Doing so allows the increase in median signal-respond RTs to be more clearly represented because individualized bins obscure this trend by collapsing over a wide range of absolute SSD values, and therefore a wide range of censoring. The resulting posterior predictive plot (Figure 2F) indicates that BEESTS-ABCD generally provides a good description of median signal-respond RTs, although it slightly underpredicts the slope of the increase and the absolute value of the highest SSD quintile. These misfits may indicate unexplained or contaminant processes at very long SSDs. However, as the absolute level of misfit is relatively small (about .025s at most), and as the general increasing trend is well described, the model appears to provide an adequate account of signal-respond RT^3^.

In sum, the BEESTS-ABCD model provides a plausible explanation of context independence violations on the ABCD task, accounting for both increasing choice accuracy rates by SSD – the hallmark of independence violations in the ABCD data – and other classic features of stop trials. We next quantify possible biases in non-parametric SSRT estimates that can occur if BEESTS-ABCD is the true data-generating process in order to gauge the impact of these biases on substantive inferences.

### Quantification of Biases in Non-Parametric SSRT

We first assessed the relationship between the parametric SSRT estimates from the BEESTS-ABCD model and those from the recommended non-parametric integration method (Verbruggen et al., 2013) that also accounts for go trial omissions (Verbruggen et al., 2019). As shown in Figure 3, the two measures shared around 60% of their variance and had a close to one-to-one relationship when the ABCD model was estimated with broad priors, but this shared variance was reduced, and the one-to-one relationship disappeared, when priors informed by the hierarchical model fit were used. The latter result reflects “shrinkage” (Efron & Morris, 1977; Gelman et al., 2013), a characteristic of hierarchical Bayesian estimation that is particularly advantageous when, as in the present case, measurement noise is high because each individual performs a relatively small number of trials. In these cases, shrinkage produces better estimates on average by pulling poorly constrained outlying estimates closer to the group mean, as is evident in the right panel of Figure 3. Regardless, these relations show that the rank ordering of participants’ parametric SSRTs was relatively well-preserved in the non-parametric estimates.

**Figure 3.**
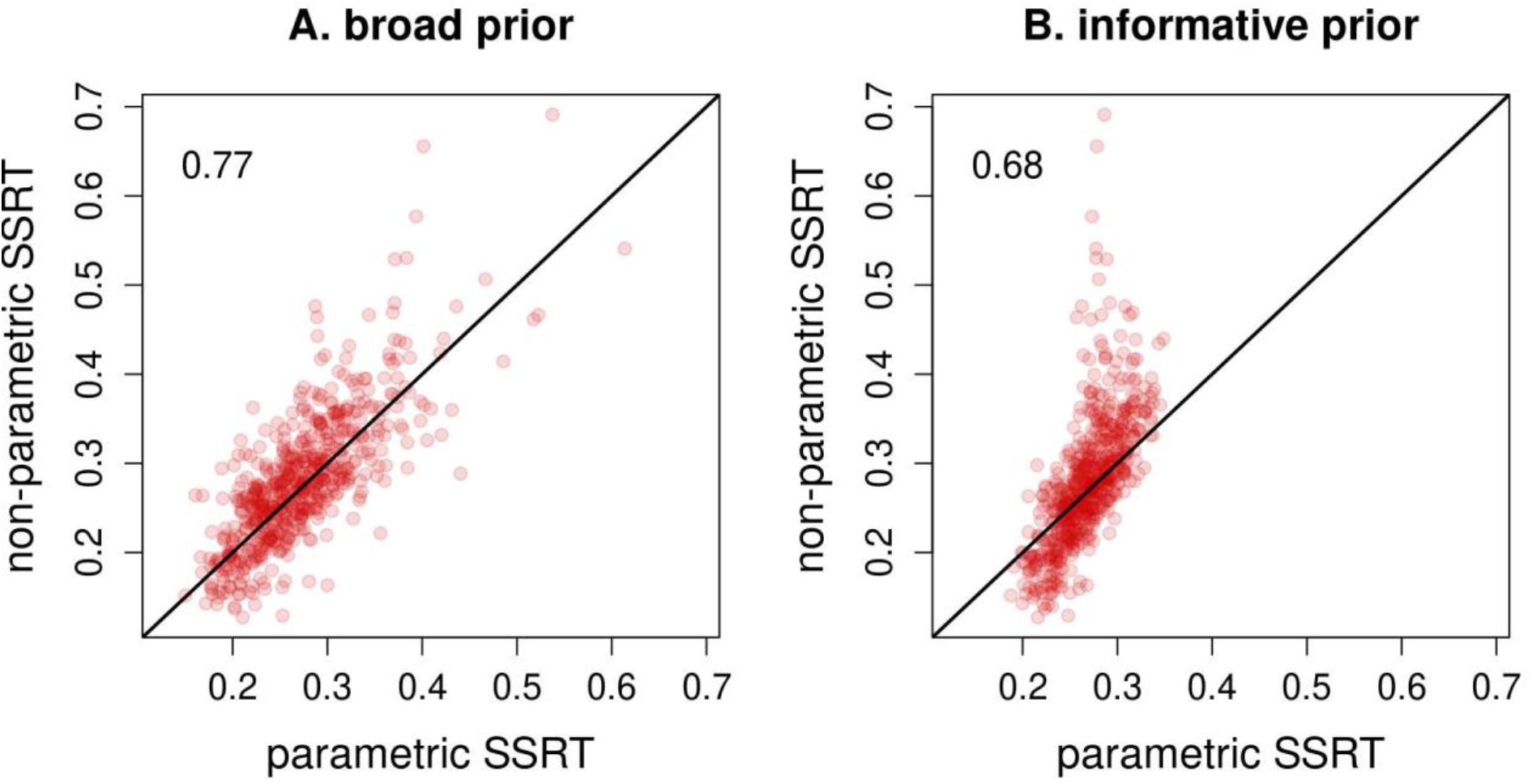
Comparison of non-parametric SSRT estimates with parametric estimates obtained from the BEESTS-ABCD model, both when the model is estimated with broad, uninformative priors (**A**) and when it is estimated with narrower priors informed by a hierarchical model fit (**B**). The black line represents where dots would fall along if the relation between the two sets of estimates was perfect. Correlation coefficients (*r*) for each relation are displayed in the top left corner of each plot.

Although the ordinal correspondence between BEESTS-ABCD and non-parametric SSRT estimates suggests that the latter can be used to study individual differences in inhibition, confounding factors that are not accounted for in non-parametric methods may nonetheless lead to misleading inferences. Indeed, we found that biases in non-parametric estimates can be consequential for inferences at both the individual and group levels. We present an example at each level to demonstrate that differences in urgency (*v*_0_) and perceptual growth rate (*g*) can be mistaken for differences in SSRT, as estimated by the non-parametric method. Note that the parameter values, and ranges over which we vary parameters, used in the simulations are representative of those found in the ABCD data^4^.

The first example shows that ignoring either parameter can lead to reversed SSRT effects when making pairwise comparisons among three individuals. Person A has a higher true parametric SSRT than the others, but roughly the same urgency as B and a similar perceptual growth rate to C. Figure 4 shows the results, where horizontal lines indicate the true parametric SSRT for each person, crosses correspond to the true urgency (left panel) and perceptual growth rate (right panel), and dots show non-parametric SSRT estimates for each simulated dataset. Varying urgency had hardly any effect on the non-parametric SSRT estimates of A, whereas a positive linear relation was found for B, creating two qualitatively different regions. When urgency was below approximately 4, A had a higher non-parametric SSRT than B, whereas this order was reversed when urgency exceeded four. By not taking urgency into account, the non-parametric SSRTs would result in incorrect conclusions in the latter region. Similarly, varying perceptual growth rate had different effects on A and C, and again created two qualitatively different regions below and above around 2.5. Non-parametric SSRT would lead to incorrect conclusions in the lower region. Both cases demonstrate that overlooking individual differences in urgency and perceptual growth rate can lead to incorrect inferences about the relative inhibitory abilities of two individuals. Unless individual differences in these factors are taken into account, as can be done using the BEESTS-ABCD the model, there is potential for the non-parametric method to identify putative differences in inhibitory ability where none exist, and even for one individual to be identified as better at inhibition than another when in fact the opposite is true.

**Figure 4.**
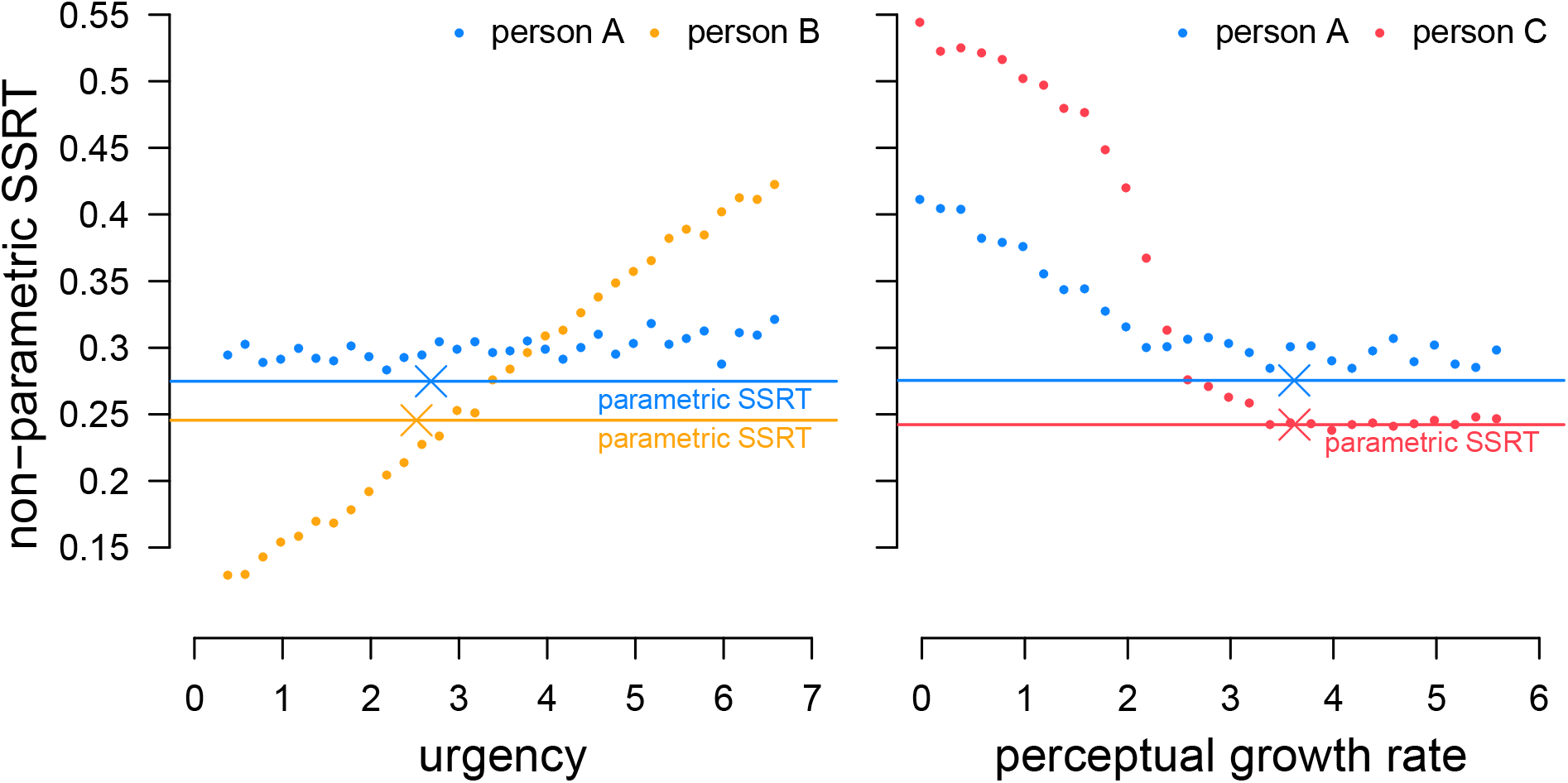
A demonstration of the influence of urgency and perceptual growth rate on the non-parametric SSRT of two individuals based on 15,000 simulated “go” and 5,000 simulated “stop” trials each. Persons A and B (left) have a similar level of urgency as indicated by the crosses, but a differing SSRT. When urgency is varied, the estimated non-parametric SSRT is affected even though this change should not lead to a different SSRT. Similarly, persons A and C (right) have a similar level of perceptual growth rate, and varying this rate also affects the estimated non-parametric SSRT. Both cases result in two regions leading to qualitatively different conclusions. In reality, person A has a higher SSRT than persons B and C, and this true parametric SSRT (horizontal lines) is independent of both urgency and perceptual growth rate. Note that with the large number of trials in these simulations BEESTS-ABCD parametric SSRT essentially are essentially perfect, and so the horizontal lines correspond to the true data generating values.

The stop-signal task is often used to identify differences in SSRT between groups or experimental conditions, or to investigate whether a covariate (e.g., ADHD symptoms or activity in a particular brain region) is related to inhibitory ability. Our second simulation shows that a covariate related to both urgency (*v_0_*) and SSRT (through the μ parameter) may cause non-parametric SSRT to provide reversed conclusions in a realistic analysis. Figure 5 shows an illustrative selection of results for 900 simulated participants. The top row shows two extreme situations in which SSRT was perfectly negatively correlated with the covariate, but there was alternately no (left plot) or a perfect positive (right plot) correlation between the covariate and urgency. Two situations with smaller, more realistic covariate correlations are presented in the bottom row.

**Figure 5.**
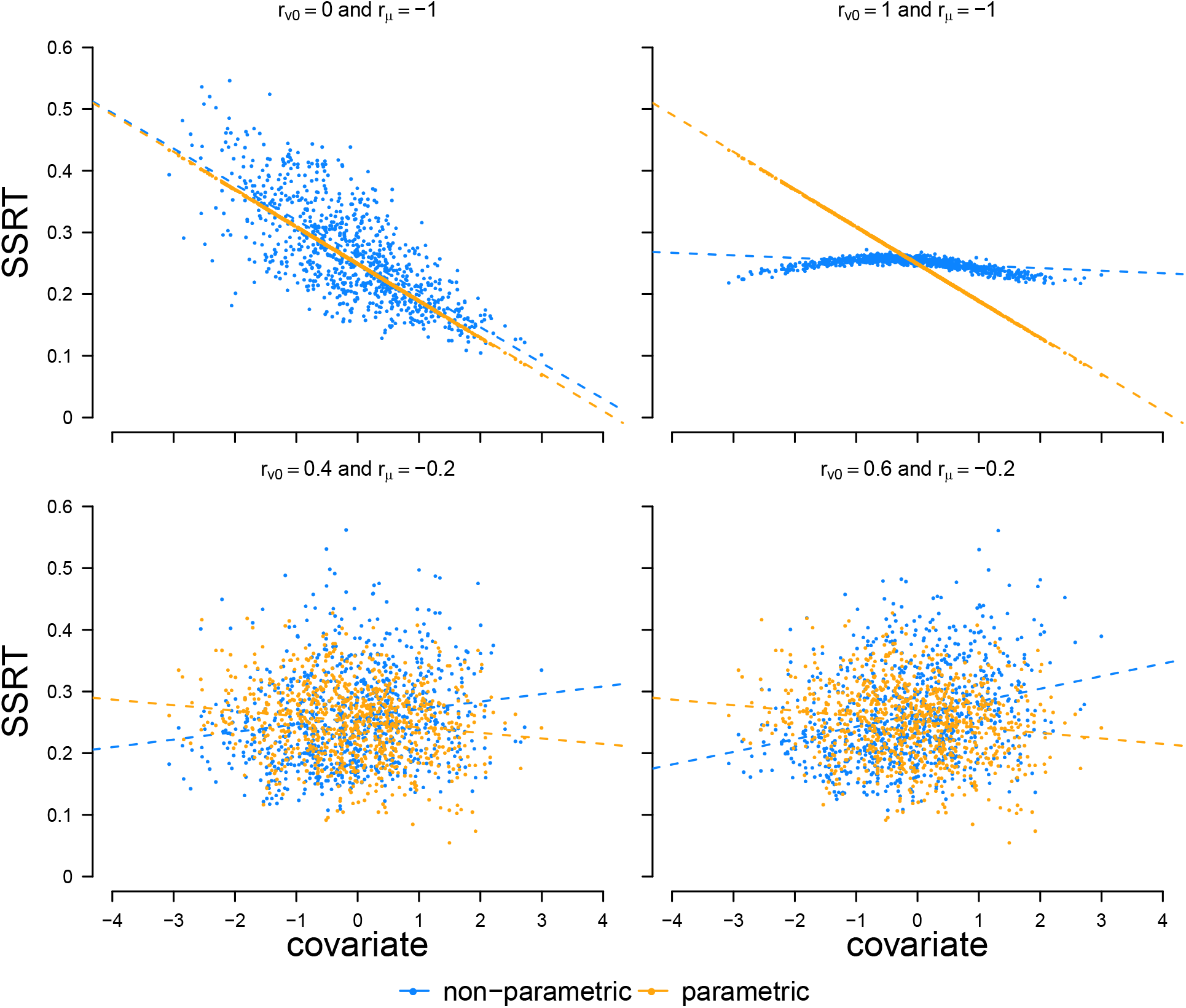
Demonstrations of covariate confounds influencing non-parametric SSRT estimates in a correlational analysis. A covariate was created that has a relation with both urgency (*v_0_*) and SSRT (through the *μ* parameter) for 900 participants, each simulated, and results obtained, in the same way as for the first example.

Both non-parametric and parametric SSRT correctly showed strong negative correlations with the covariate when only SSRT was perfectly related to the covariate (top-left). However, their results clearly diverged when urgency also had a perfect positive correlation with the covariate (top-right). In this case, parametric SSRT had the expected strong negative relation with the covariate, while non-parametric SSRT estimates hardly varied over different levels of the covariate. Non-parametric SSRT estimates failed to pick up the negative relation between the covariate and true SSRT because of the counter-balancing effect of the positive urgency correlation on non-parametric SSRT. The bottom row of Figure 5 demonstrates that smaller covariate correlations, closer to those expected in empirical data, can also result in reversed effects. In both scenarios, BEESTS-ABCD accurately detected a small *negative* relation between the covariate and SSRT. From the non-parametric estimates, however, we would conclude that there is a *positive* relation between the covariate and SSRT because the positive correlation with urgency reverses this trend. These results provide a few examples of the many possible ways inhibition-related covariate effects (or the absence of such effects) can be confounded by urgency effects. Growth rate effects can produce similar confounding. Therefore, failing to take different levels of context-independence violation into account can, in realistic scenarios, lead to qualitatively wrong conclusions despite the ordinal correspondence between true and non-parametric SSRT.

### The ABCD Stop-Signal Model as a Measurement Model

Given that non-parametric estimates of SSRT may cause misleading inferences, it is natural to ask whether the BEESTS-ABCD model can be used to avoid such problems with data from the ABCD design. Even though the model provides a good description of empirical trends in the ABCD data, this feature alone does not necessarily mean that it can serve as *measurement model*. Measurement models must have a one-to-one mapping between data-generating model parameters and parameter estimates that can be obtained from a realistic design (Matzke et al., 2020). This can be checked via simulation in a parameter-recovery study (Heathcote et al., 2015).

We conducted parameter-recovery studies using parameter estimates from our subsample to generate simulated individual data sets with same trial numbers and staircase algorithm as the empirical ABCD stop-signal data. We then used the same procedures as applied to the empirical data to estimate BEESTS-ABCD parameters from the simulated data. As priors can impact parameter recovery in Bayesian estimation, we repeated this procedure with both the informed and broad priors used with the empirical data. Figures 6 and 7 display scatterplots of the relations between the data-generating and estimated parameters, associated correlation coefficients (*r*), and posterior “coverage” (*c*), the proportion of data-generating parameter values that fall within the 95% posterior credible intervals provided by the Bayesian estimation procedure. If the coverage proportions are close to the nominal 95% value it indicates that estimation is *calibrated* in terms of uncertainty.

**Figure 6.**
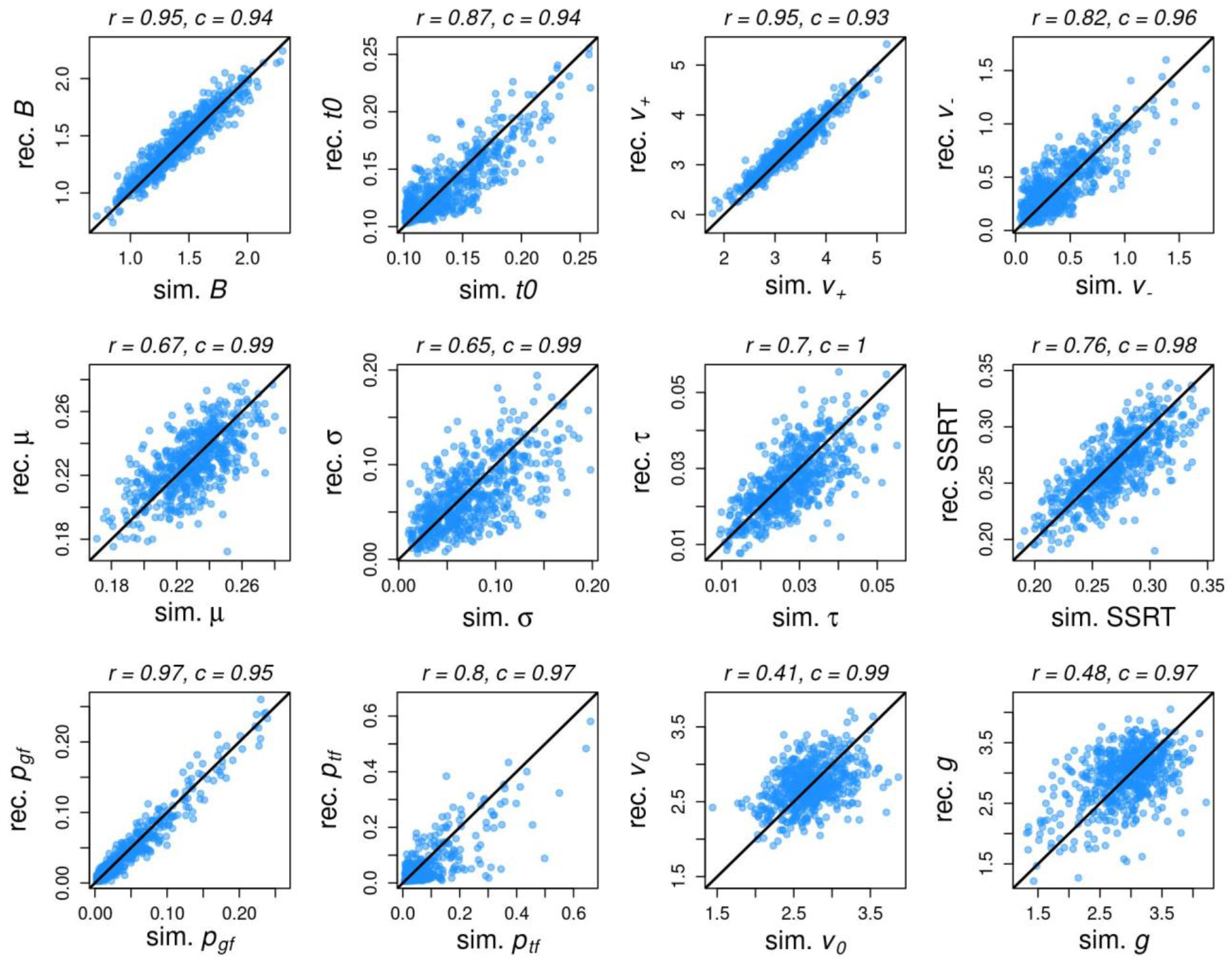
Results from the parameter recovery study in which informed priors were used to estimate parameters (both the initial parameter values estimated from empirical data and the parameter values recovered from simulated data). Scatterplots illustrate the relations between the simulated (“sim.”) and recovered (“rec.”) parameter values as compared to the diagonal solid line indicating perfect recovery. Numbers above each plot report the correlation coefficient (*r*) for each relation and the posterior coverage proportions (*c*) for each parameter, which indicate the proportion of data-generating parameter values that fall within the 95% posterior credible interval for the parameters recovered from the generated data.

**Figure 7.**
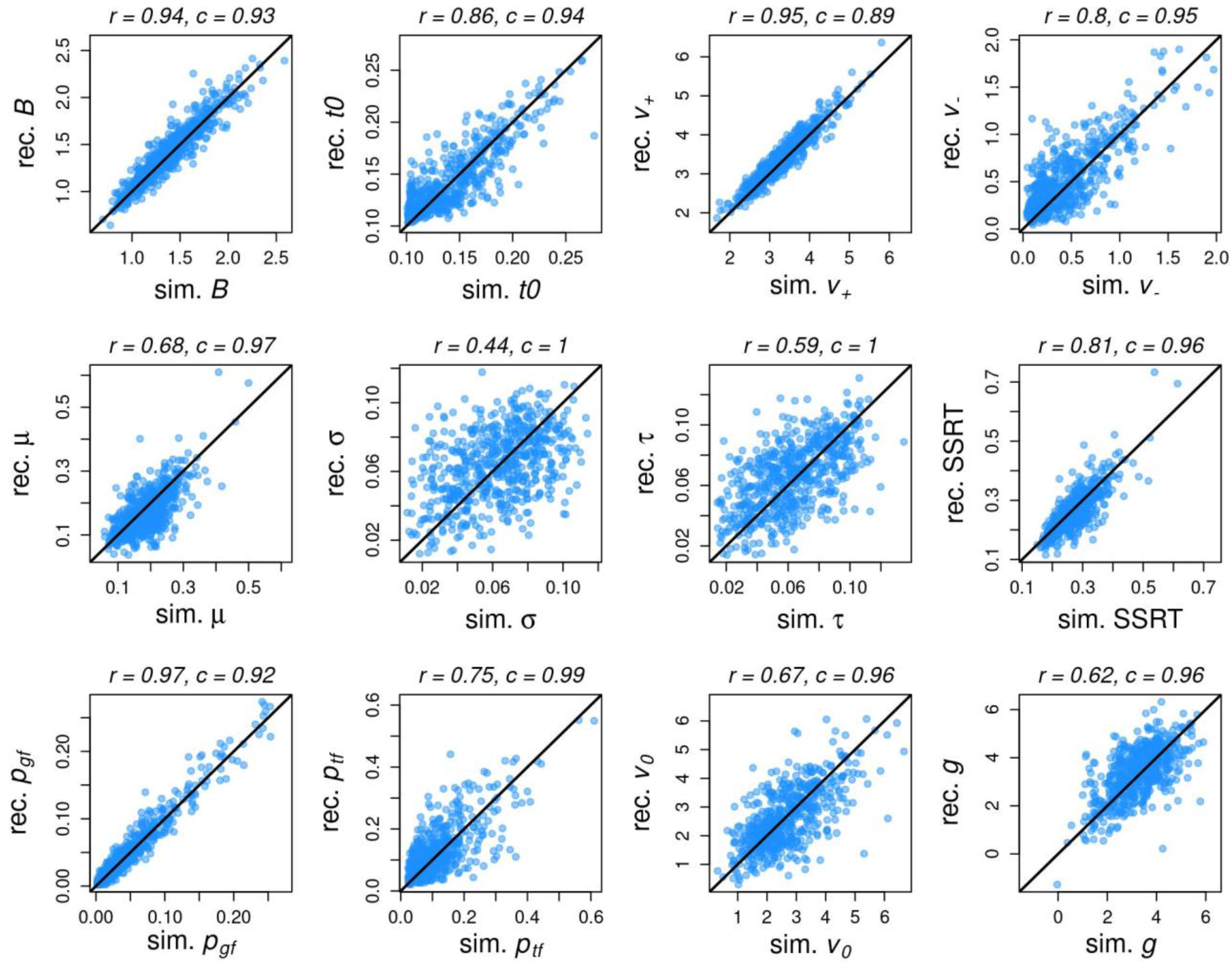
Results from the parameter recovery study in which broad priors were used to estimate parameters (both the initial parameter values estimated from empirical data and the parameter values recovered from simulated data). Scatterplots illustrate the relations between the simulated (“sim.”) and recovered (“rec.”) parameter values as compared to the solid diagonal line indicating perfect recovery. Numbers above each plot report the correlation coefficient (*r*) for each relation and the posterior coverage proportions (*c*) for each parameter, which indicate the proportion of data-generating parameter values that fall within the 95% posterior credible interval for the parameters recovered from the generated data.

As the points in Figures 6 and 7 do not fall systematically higher or lower than the line indicating perfect recovery for any parameter, there is little evidence of estimation bias^5^. Coverage of the recovered 95% credible intervals was nominal, or very close to nominal, in all cases, suggesting that inference based on credible intervals will be well calibrated. The parameters which characterize the go process, and its probability of being triggered (*v*_+_, *v*_-_, *B*, *t*_0_, and *p*_gf_), are all recovered quite precisely (*r*=0.80-0.97). Parameters that characterize the stop process and its probability of being triggered (*μ*, *σ*, *τ*, *p*_tf_) are less precise, but generally acceptable, except for *σ* in the broad prior estimates, which is notably poor. Less precision for stop, relative to go, parameters is expected because stop trials are less frequent, and this loss of precision also occurs in standard paradigms for the BEESTS-trigger-failure model. Crucially, the BEESTS-ABCD model recovers mean SSRT and trigger failure values well (*r*=0.75-0.81). Given that 60 stop trials is barely above the minimal guideline for using non-parametric estimates in the standard stop-signal paradigm (Verbruggen et al., 2019) this represents excellent performance and suggests that the two key stop-related parameters estimated from BEESTS-ABCD can be used effectively in applied research.

Parameters for urgency (*v*_0_) and perceptual growth (*g*), which similarly depend on the sparse stop trial data, displayed poor recovery. However, as they are more likely to be considered nuisance parameters than of substantive interest, and as there were no apparent biases in estimates of these parameters, this is unlikely to limit applications of the model. Furthermore, the coverage values indicate that, even if a subset of parameters’ point estimates are not precisely recovered, the posterior distributions accurately reflect the uncertainty in these estimates and can therefore inform appropriately tentative inferences.

## Discussion

We proposed a new cognitive process model, BEESTS-ABCD, that accounts for context independence violations in the ABCD Study stop-signal task that are due to a design feature in which the visual stop signal replaces the go choice stimulus, effectively limiting the information participants need to make a choice. By combining elements of prior parametric models of the stop-signal task (Logan et al., 2014; Matzke et al., 2013; Matzke, Love, et al., 2017) with accounts of masking effects on the processing of brief visual stimuli (Ratcliff & Rouder, 2000; P. L. Smith & Ratcliff, 2009; P. L. Smith & Sewell, 2013), the model assumes that the ABCD design feature impacts the quality of evidence used for discrimination of choice options. The model provides an excellent account of the pattern of increasing choice accuracy over SSDs, which is the “smoking gun” that the task violates context independence (Bissett et al., 2021), and a good account of other more standard features of stop trial performance. We found that the rank ordering of parametric SSRT values from this model is generally well-preserved in non-parametric SSRT estimates, which do not account for the context independence violations. However, we also show in simulation studies that confounding factors can lead non-parametric estimates to reverse the ordering of SSRT differences at both the individual and group levels. Critically, parameter recovery studies demonstrate that, when fit to data from the ABCD design, BEESTS-ABCD can be used to avoid these problems by reliably and validly measuring SSRT, trigger failures and other processes of interest.

Given the unprecedented scientific opportunity afforded by the ABCD Study and the importance of unbiased SSRT estimates for researching inhibitory ability, these findings have several significant implications. First, they suggest that prospective ABCD data collection can continue without changing the task design, ensuring longitudinal comparability of the behavioral and fMRI data between data collection waves. SSRT measurement issues related to the violation are manageable using BEESTS-ABCD, and the practical difficulties of using the model are relatively minor compared to the problems introduced by breaking longitudinal comparability. Furthermore, even if alternate models that explain the context-independence violation with different processes are proposed and supported, we believe our results provide a general demonstration that cognitive-model-based solutions can provide an effective way to overcome limitations related to the ABCD design.^6^

The implications of our work for analyses of already-collected ABCD data are nuanced. As we found that non-parametric SSRT estimates calculated using recommended best practices (Verbruggen et al., 2019) generally preserved the rank ordering, if not the absolute values, of participants’ SSRT, it is possible that inferences based on the non-parametric estimates may not be misleading in many situations. However, we also showed that non-parametric estimates can lead to incorrect (including reversed) inferences when a parameter that explains the context independence violations is confounded with a covariate of interest. It seems plausible that such confounding might occur in practice; for example, urgency has been linked to impulsivity, depression and Parkinson’s Disease (Carland et al., 2019). Rather than taking the chance of assuming that such confounding is not present, we recommend researchers to use parametric measurement models, such as the one we propose, that account for the context-independence violations evident in the ABCD data.

We note that this trade-off between the precision of cognitive process modeling and the ease of using non-parametric SSRT estimates is not unique to the ABCD task. As outlined in the introduction, trigger failures in standard designs cannot be easily accommodated using non-parametric methods and have already been shown to bias SSRT estimates (Matzke, Love, et al., 2017) and distort substantive conclusions (Matzke, Hughes, et al., 2017; Weigard et al., 2019) if ignored. Indeed, current consensus recommendations for estimating SSRT from the stop-signal task (Verbruggen et al., 2019) acknowledge that cognitive process models, despite being difficult for researchers who are less technically adept to implement, provide less biased estimates of SSRT relative to even the best non-parametric methods. Our findings suggest parallel recommendations for analysis of the ABCD task; whenever possible, cognitive process models such as BEESTS-ABCD should be used to estimate SSRT and trigger failure while accounting for ABCD-specific context-independence violations.

To reduce technical barriers to the adoption of BEESTS-ABCD, we have shared, on the Open Science Framework (osf.io/2h8a7), the code we used to specify and fit the model within Dynamic Models of Choice (DMC), a free set of R functions for Bayesian estimation of evidence accumulation models (Heathcote et al., 2019). Researchers can freely use our code to estimate parameters of our model for any subset of ABCD participants they wish using identical procedures to those implemented in the current study. However, a centralized effort to estimate parameters from our model, for the whole sample, that can be included in future ABCD Study data releases would be ideal. We would be look forward to contributing to such an effort if the ABCD community agrees that doing so is appropriate.

We used priors generated from a hierarchical model fit to an independent subsample of ABCD participants to inform individual-level estimation because this method provides the key benefit of a fully hierarchical approach (shrinkage informed by group distributions) without some of its drawbacks. Specifically, fitting hierarchical models to very large data sets is demanding in terms of computational resources and technical expertise, although it has been done for other large-scale projects (PISA, 2018). We believe developing such large-scale hierarchical cognitive-process models is a worthwhile aim for not only the stop-signal task, but also the other tasks used in the ABCD project. However, here we focused on individual-analysis methods that require only a modern multi-core PC, so that the benefits of BEESTS-ABCD are more immediately available. Another drawback is that individual-level estimates from hierarchical models are unsuited for follow-up frequentist or Bayesian inferential methods (Boehm et al., 2018). “Plausible values” analyses have been proposed as a solution for bivariate correlations with hierarchical cognitive-model parameters (Ly et al., 2017), but until this approach is extended to the multi-level modeling, structural equation modeling, and multivariate prediction methods commonly used with ABCD data (which is another critical area for future work), individual-level estimation with informed priors provides a good alternative solution.

Further work may also be required to develop a more detailed understanding of the mechanisms mediating the brief-presentation effects in the ABCD design. There is a hint in some elements of apparent BEESTS-ABCD model misfit, such as underprediction of signal-respond RTs at long SSDs, that a more fine-grained account may require the model to be extended or a different approach developed, perhaps based on the preliminary modeling frameworks outlined by Bissett et al. (2021). However, a key strength of the current model is its parsimony; it provides a precise account of choice accuracy across SSDs and of inhibition functions by only adding two parameters to account for context independence violations. We focused on this relatively simple model because we suspect that data from the ABCD design is not suitable for developing more complex models of individual performance, as a much larger number of trials is typically necessary to estimate parameters from such models (P. L. Smith & Little, 2018). Even if such a refined model were developed, then following the aphorism summarizing the message of Box (Box, 1976) that “all models are wrong, but some are useful”, it would still need to be established whether it provides a markedly better measurement model for the ABCD design than BEESTS-ABCD does.

In summary, we propose a cognitive-process model that explains the impact of context-independence violations on the ABCD Study’s stop-signal task and, in doing so, accounts for key trends in the ABCD data. We show that failing to account for context-independence violations can produce misleading inferences, and that the proposed model provides a practical remedy, enabling unbiased and reliable estimation of SSRT and other key process parameters that contribute to task performance. We suggest that the model can advance ABCD Study research efforts by improving the measurement of inhibition and other cognitive processes (e.g., trigger failure and choice evidence accumulation) with existing ABCD stop-signal task data and by allowing for longitudinal data collection to move forward without a need for major changes to the task design. More broadly, this work highlights the critical strengths of a cognitive-process-modeling approach for increasing the precision of both theories and measures of neurocognitive

phenomena.

## Materials and Methods

### Participants, Inclusion Criteria and Subsample Selection

The ABCD Study is a multi-site collaboration that has recruited a diverse sample of 11,875 children between the ages of 9 and 10 from 21 study sites across the United States. Details of the general study design, recruitment procedure, and fMRI protocol (including the stop-signal task design) are available elsewhere (Casey et al., 2018; Garavan et al., 2018). The ABCD data repository, which is openly available via the National Institute of Mental Health Data Archive (NDA), grows and changes over time. As the Bayesian estimation procedures that we used to fit the proposed cognitive process model to data from the ABCD stop-signal task are computationally intensive, fitting the model to the full sample during model development and testing would be unnecessarily time intensive. Therefore, we randomly selected a subset of 900 individuals (300 for the derivation of informed priors and 600 for the primary analyses) whose baseline session stop-signal data met our performance-based inclusion criteria. We selected this subset from the set of individuals whose baseline stop-signal data were included in the ABCD Annual Release 1.1 dataset (*n*=4,521, NDA Study 576, DOI 10.15154/1412097, available at: https://ndar.nih.gov/study.html?id=576).

Of the 4,521 individuals in Release 1.1, we found that 3,436 had complete behavioral data available from the stop-signal task that met basic validity checks (two imaging runs, 360 recorded trials, and participant responses detected for one or more go trials). We then applied the following performance exclusion criteria to ensure that individuals included in our analyses were adequately engaged in the task and had behavioral data that were broadly consistent with race model assumptions:

a. Choice accuracy rate <.55, indicating inattention or misunderstanding of the choice task (1.9% of the sample).
b. Excessive rate of omission (>.25) on go trials (4.2% of the sample).
c. Low overall rate of successful inhibition (<.25) on stop trials (1.2% of the sample).
d. Mean RT on stop trials was greater than mean RT on go trials, indicating gross violations of race model assumptions (6.0% of the sample).
e. Following Weigard et al. (2019), we assessed whether individuals’ mean RT changed substantially (>.500s) over the course of the task, which would violate race model assumptions, by fitting a linear model in which RT on a given trial was predicted by trial number (1-360). The resulting coefficient was multiplied by 360 to estimate the total RT change over the course of the task, and individuals with an absolute RT change value of 0.5s were excluded (0.2% of the sample).
f. The race model cannot explain inhibition functions in which probability of responding decreases as SSDs get longer. To evaluate whether individuals displayed evidence of decreasing inhibition functions, we fit a logistic regression model to individuals’ stop trial data in which probability of responding was predicted by SSD. Individuals with negative regression coefficients, indicating that overall probability of responding decreased as SSDs grew longer, were excluded (5.0% of the sample).
g. Presence of the rare tracking algorithm glitch noted by Bissett et al. (2021; “Issue 3”), which incorrectly causes SSDs to remain stuck at the minimum value of 0s (2.1% of the sample).
h. Presence of SSDs >0.7s, which, as noted by Bissett et al. (2021; “Issue 4”) causes the visual stop signal presentation time to be truncated (3.9% of the sample).

Simultaneous application of all the above criteria led to the exclusion of 16.8% of the initial sample of 3,436, leaving 2,859 participants available for modeling analyses. Out of this remaining sample, we sought to randomly select a subsample of non-sibling participants from across multiple data collection sites. We first determined, for each data collection site, how many individual family identification numbers (family IDs) had stop-signal data available for modeling from at least one child. We then selected the 6 data collection sites with the greatest number of available family IDs and randomly sampled (without replacement) 150 family IDs from each of these 6 sites. We included single children from each family for which only one child was available and randomly sampled a sibling from each family for which multiple children’s data were available. This led to a subsample of 900 participants from independent families (150 per site) that was further split into a 300 participant (50 per site) subsample included in hierarchical model fits to estimate parameter priors and a 600 participant (100 per site) subsample included in the main analyses. Table 2 displays basic demographic characteristics of the 900 participants selected for analysis.

**Table 2.** Demographic characteristics of the 900 ABCD participants meeting performance-based inclusion criteria whose data was randomly selected for modeling analyses. k = $1,000 (United States) of annual income; GED = General Educational Development test passed, equivalent to a High School education.

| **Demographic** | **Category** | ***n*** | **%** |
| --- | --- | --- | --- |
| Sex | *Female* | 422 | 46.9 |
|  | *Male* | 478 | 53.1 |
| Household Income | *<50k* | 270 | 30.0 |
|  | *>=50k & <100k* | 286 | 31.8 |
|  | *>=100k* | 266 | 29.6 |
| Highest Parental Education | *<High School* | 42 | 4.7 |
|  | *High School / GED* | 78 | 8.7 |
|  | *Some College* | 260 | 28.9 |
|  | *Bachelor* | 214 | 23.8 |
|  | *Post-Graduate* | 300 | 33.3 |
| Household Marital Status | *Married* | 594 | 66.0 |
|  | *Not married* | 299 | 33.2 |
| Race | *White* | 602 | 66.9 |
|  | *Black* | 73 | 8.1 |
|  | *Asian* | 17 | 1.9 |
|  | *Other/Mixed* | 184 | 20.4 |
| Ethnicity | *Hispanic* | 299 | 33.2 |
|  | *Not Hispanic* | 591 | 65.7 |

### Model Estimation

Details of model parameterization are outlined above in the main text. We implemented the 11-parameter BEESTS-ABCD model within Dynamic Models of Choice (DMC), a free set of R functions for Bayesian estimation and simulation of evidence accumulation and stop-signal task models (Heathcote et al., 2019). Following previous work (Matzke, Hughes, et al., 2017; Weigard et al., 2019), parameters for the probability of trigger failure (*p*_tf_) and “go” failure (*p*_gf_) were projected onto the real line during model estimation using a probit transformation and later transformed back to the natural scale for interpretability (e.g., in Figures 6 and 7, above). Before all modeling analyses, trials with RTs <0.15s were excluded as fast guesses and trials with RTs >1.5s were excluded as abnormally slow responses (these exclusions removed data from <1% of trials). The model was fit, using individual-level Bayesian estimation (specific procedures described below), to the main 600-participant subsample twice: once with broad, relatively uninformative priors and a second time with informative priors derived from hierarchical model fits to the independent subsample of 300 participants. All priors were truncated normal distributions with the bounds and distributional parameters listed in Table 3.

**Table 3.** Prior probability distributions used for Bayesian estimation of the ABCD stop-signal model parameters. All priors were truncated normal distributions with the bounds, location parameters and scale parameters listed here. Location and scale parameters for the “Broad” prior distributions were selected a priori. Location and scale parameters for the “Informative” prior distributions were derived from a hierarchical Bayesian version of the ABCD stop-signal model that was fit to a subset of 300 ABCD participants using procedures described in the text. Note that priors for the *p*_gf_ and *p*_tf_ parameters are represented on the probit scale.

| Parameter | *Bounds* |  | *Broad*<br>*Prior Distribution* |  | *Informative*<br>*Prior Distribution* |  |
| --- | --- | --- | --- | --- | --- | --- |
|  | lower | upper | location | scale | location | scale |
| $t_0$ | 0.100 | 1.000 | 0.150 | 0.100 | 0.027 | 0.068 |
| $B$ | 0.000 | ∞ | 1.000 | 1.000 | 1.444 | 0.308 |
| $v_+$ | 0.000 | ∞ | 3.000 | 2.000 | 3.334 | 0.550 |
| $v_-$ | 0.000 | ∞ | 1.000 | 2.000 | -0.045 | 0.525 |
| $v_0$ | 0.000 | ∞ | 2.000 | 2.000 | 2.732 | 0.634 |
| $g$ | $-\infty$ | ∞ | 3.000 | 2.000 | 2.958 | 0.862 |
| $\mu$ | 0.000 | 2.000 | 0.500 | 0.300 | 0.226 | 0.036 |
| $\sigma$ | 0.000 | 0.500 | 0.050 | 0.100 | 0.012 | 0.095 |
| $\tau$ | 0.000 | 0.500 | 0.100 | 0.100 | 0.014 | 0.032 |
| $p_{gf}$ | $-\infty$ | ∞ | 0.000 | 1.000 | -1.944 | 0.517 |
| $p_{tf}$ | $-\infty$ | ∞ | 0.000 | 1.000 | -1.892 | 0.764 |

The hierarchical modeling approach that was used to obtain informative priors treats individual-level parameters as random effects described by group-level truncated normal distributions that are defined by a location hyperparameter and a scale hyperparameter. Priors for the location hyperparameters were the same broad prior distributions displayed in Table 2 while priors for all scale hyperparameters consisted of exponential distributions with a scale of 1. Following the estimation of posterior distributions from the hierarchical model for the 300-person subsample, we collapsed the individual-level posterior samples across all 300 individuals into a single vector for each model parameter. We then fit a truncated normal distribution (with bounds for each parameter specified in Table 2) to the distribution of samples in each parameter vector using maximum likelihood. The resulting location and scale parameters of the fitted truncated normal distributions (Table 2) were used to specify informative priors for the fits to the remaining 600 participants.

All hierarchical and individual-level Bayesian parameter estimation procedures used the differential-evolution Markov chain Monte Carlo (DE-MCMC) algorithm to sample from the posterior, as this sampling method is more efficient for the estimation of evidence accumulation models and similar models that tend to have correlated parameters (Turner et al., 2013). These simulations included a number of chains that, by DMC default, was three times the parameters in the model (e.g., 33 chains for the 11-parameter ABCD stop-signal model). Each simulation featured an initial burn-in period that included a migration (Turner et al., 2013) step (with 5% probability in individual-level fits and 2.5% probability in hierarchical fits) and lasted until no chains were repeatedly “stuck” in one location for many iterations, as determined by an automated function in DMC with default settings. Next, a second burn-in period was started with migration turned off and lasted until chains for all parameters had converged on a stable posterior. Final convergence was defined as values of <1.10 for the Gelman-Rubin diagnostic statistic (Gelman et al., 1992), and was corroborated via visual inspection of the chains (Lee & Wagenmakers, 2014). Following initial sampling, mean SSRT samples were computed by simulating 10,000 finishing times from ex-Gaussian distributions defined by each set of *μ, σ,* and *τ* samples and taking the mean of finishing times over the .05s lower bound.

The same procedures outlined above were also used to estimate a “go independence” model for comparison of its ability to describe choice accuracy on stop trials (shown in Figure 2C) with the ability of the ABCD stop-signal model (Figure 2B). This model had an identical structure to the ABCD stop-signal model except that it assumed that the go process across stop trials of all SSDs had identical accumulation rates to the go process on go trials (i.e., *v*_0_ and *g* were not estimated). The fits shown in Figure 2C were from individual-level Bayesian estimation of this model that, similar to the fits of the main ABCD stop-signal model shown throughout Figure 2, used informative priors derived from a hierarchical fit of the “go independence” model to the separate subset of 300 participants.

Posterior predictions were generated by drawing 100 samples from the 11-parameter joint posterior and simulating data predicted by the model for each participant and each of the 100 sets of parameter samples. The predicted data for each of the 100 sample sets were averaged across participants within each set to obtain summary values of interest (e.g., average accuracy rates and probability of inhibition at specific SSDs). The medians and 99% credible intervals (CIs) of these group-average predictions were then estimated with 100 draws of posterior samples and plotted against the empirical group-average values in Figure 2.

### Comparisons to Non-parametric Methods

We set up two simulation studies to demonstrate how the BEESTS-ABCD model SSRTs and non-parametric estimates can differ when not accounting for context-independence-violation. The first simulation is set up at the individual level, and the second at the group level.

For the individual example, we picked three individuals (persons A, B, and C) from a subset of ABCD data, and used their parameters obtained from fitting BEESTS-ABCD to their data. The parameter estimates used here and in the group example were obtained using broad priors, but similar results are possible from estimates obtained using informed priors. Person A had a higher parametric SSRT than the others, but roughly the same urgency as B and similar perceptual growth rate to C. To demonstrate the potential effect of urgency, we used the parameter estimates of person A and B to simulate new data sets with a large number of trials (15,000 simulated “go” and 5,000 simulated “stop” trials) so that results are clear. All parameters, except urgency, were fixed to their estimates for both persons. Urgency was varied over a realistic range (i.e., within the range found in our fits to ABCD data). For each level of urgency, we simulated new datasets for each person while keeping all other parameters constant. We followed a similar approach to inspect the potential effect of perceptual growth rate. Instead of varying urgency, we varied the perceptual rate and compute non-parametric SSRT in each data set. For each generated dataset, we computed non-parametric SSRT estimates and compared these estimates to the BEESTS-ABCD SSRTs used to generate the data.

For the group example, we generated new datasets in which both urgency and SSRT were correlated with a covariate to varying degrees (*r* = −1 to *r* = 1 in steps of .2). To ensure that all data-generating parameters had realistic values, we set (i) the mean and standard deviation of *v_0_* and to the values found in the BEESTS-ABCD fits to a subset of ABCD data; (ii) all other data-generating parameters to the median of estimates from ABCD data; and (iii) perceptual growth rate to a slightly lower, but still realistic, value. From these parameter settings, we simulated new datasets with 900 participants in the same way as for the first example, for each combination of correlations between the covariate and *v_0_*, and the covariate and μ. Again, non-parametric SSRT estimates were computed and compared to the BEESTS-ABCD SSRTs used to generate the data.

### Parameter Recovery Studies

We conducted two different parameter recovery studies based on parameter estimates from all participants in the main 600-person subsample. The first study used the posterior medians of individual-level Bayesian parameter estimates obtained under the informative priors derived from the earlier hierarchical model fit. The second study used the posterior medians of individual-level Bayesian parameter estimates obtained under the broad, relatively uninformative priors. In both recovery studies, the 600 sets of posterior median parameter estimates were used to simulate 600 individual-level data sets that each had identical trial numbers (300 “go”, 60 “stop”) and an identical staircase tracking procedure to the empirical ABCD data. Next, individual-level Bayesian estimation was used to estimate parameters from the simulated data sets, with the informative priors and broad priors used in the first and second recovery studies, respectively (i.e., the priors used to fit the original empirical data and the corresponding simulated data were identical in each case). Posterior medians of the resulting parameter estimates were then compared with the data-generating parameter values in each case (see Figures 6 and 7). Recovered posterior medians of mean SSRT were computed using the same simulation procedure, outlined above, that was used to compute mean SSRT for the empirical data.

We also completed more intensive parameter recovery studies, similar to those conducted in Matzke et al. (2020), to better assess the possibility of biases or second modes within the parameter space. We randomly drew participants who had *B*, *t*_0_, *p*_tf_, *μ*, *τ*, *v*_0_ and *g* posterior median parameter estimates within each of the 4 quartiles that characterized individual variation in each parameter, leading to 28 unique parameter sets. We then used each of these parameter sets to simulate 200 individual-level data sets with identical features to the ABCD stop-signal task and re-estimated parameters from each of these data sets using individual-level Bayesian estimation. Plots comparing the values of each data-generating model parameter to posterior median parameter estimates obtained from the 200 simulated data sets, which are available for download on our OSF project page (osf.io/2h8a7/), indicated little evidence for substantial biases or for second modes in the parameter space.

## Supplemental Materials

### 0-Second SSD Responding Uptick Noted by Bissett et al. (2021)

As stated in the Results section, Bissett et al. (2021) show that when stop trials are binned by specific SSDs there is a gradual decrease in the average probability of responding as SSDs get shorter, as predicted by the race model, but also a slight uptick in responding on 0s SSD trials relative to other short-SSD trials, which is unexpected given the assumptions of the race model. We did not see any evidence of this uptick in our initial analyses because of our strategy of plotting inhibition functions while using the individual-level binning procedure outlined in the Results section (and implemented in Figure 2D), which has the advantage of accounting for individual differences in inhibitory performance. However, after seeing the 0s SSD uptick highlighted in a revised version of the Bissett et al. (2021) preprint manuscript, we generated posterior predictive plots of the inhibition function with trials binned by individual SSDs and found that the 600 participants in our analytic subsample also displayed this group-average pattern. Curiously, we found that the BEESTS-ABCD model we propose can account for this uptick on 0s SSD trials when it is fit with broad, uninformative priors (Supplemental Figure 1A), but has more difficulty accounting for it when the model is fit with informative priors derived from a hierarchical model (Supplemental Figure 1B). This was particularly surprising given that we typically found improvements in the model’s fit to most other group-average trends in posterior predictive plots when informative priors, relative to broad priors, were used.

The sensitivity of this group-average trend to priors, combined with the fact that the uptick was not apparent in inhibition function plots that used our individual-level binning procedure, led us to suspect that the apparent increase in responding on 0s SSD trials may not represent a mechanistic pattern occurring at the level of individual subjects. Rather, we thought the uptick may be an artifact that appears when data are aggregated by SSD across individuals of very different levels of performance because the participants with the worst performance have both: i) the most 0s SSD trials and ii) the highest probability of responding on those trials. To assess the validity of this hypothesis, Supplemental Figure 1C compares the group-average trend (black dots) with the average response probabilities of three subgroups stratified by their number of 0s SSD trials: 1) subjects with no 0s SSD trials (blue line), 2) subjects with “few” (1-9) 0s SSD trials (orange line), and 3) subjects with “many” (>=10) 0s SSD trials (red line). The “many” subgroup represents only 10% of the sample but accounts for the majority (61%) of the 0s SSD trials. The “few” subgroup represents about a third (34%) of the sample and only accounts for a minority (39%) of the 0s SSD trials. These proportions suggest that the “many” subgroup, which has the worst inhibitory performance, has a disproportionate influence on the group average for responding at 0s SSDs. As expected, the comparison plot shows that there is no evidence for a 0s SSD uptick when the three subgroups are considered separately. Furthermore, the plot demonstrates that the apparent uptick is caused by the fact that individuals with no 0s SSD trials (the majority of the sample) have an inhibition function that decreases to 0 at .05s SSD trials (as expected due to tracking algorithm) but individuals with 0s SSD trials tend to have high response rates on these trials because of their poorer inhibitory performance.

This led us to conclude that the uptick noted by Bissett et al. (2021) is an artifact of group averaging rather than a mechanistic feature of performance at the individual level that must be accounted for by a process model. The BEESTS-ABCD model is able to account for this group average trend when it is unconstrained by priors, but likely has more difficulty doing so when constrained by hierarchical-model-informed priors because of “hierarchical shrinkage” (Gelman et al., 2013), which prevents overfitting by pulling parameter estimates of the worst performing participants (i.e., those with the greatest number of 0s SSD trials) closer to the group mean.

---

**Supplemental Figure 1.**
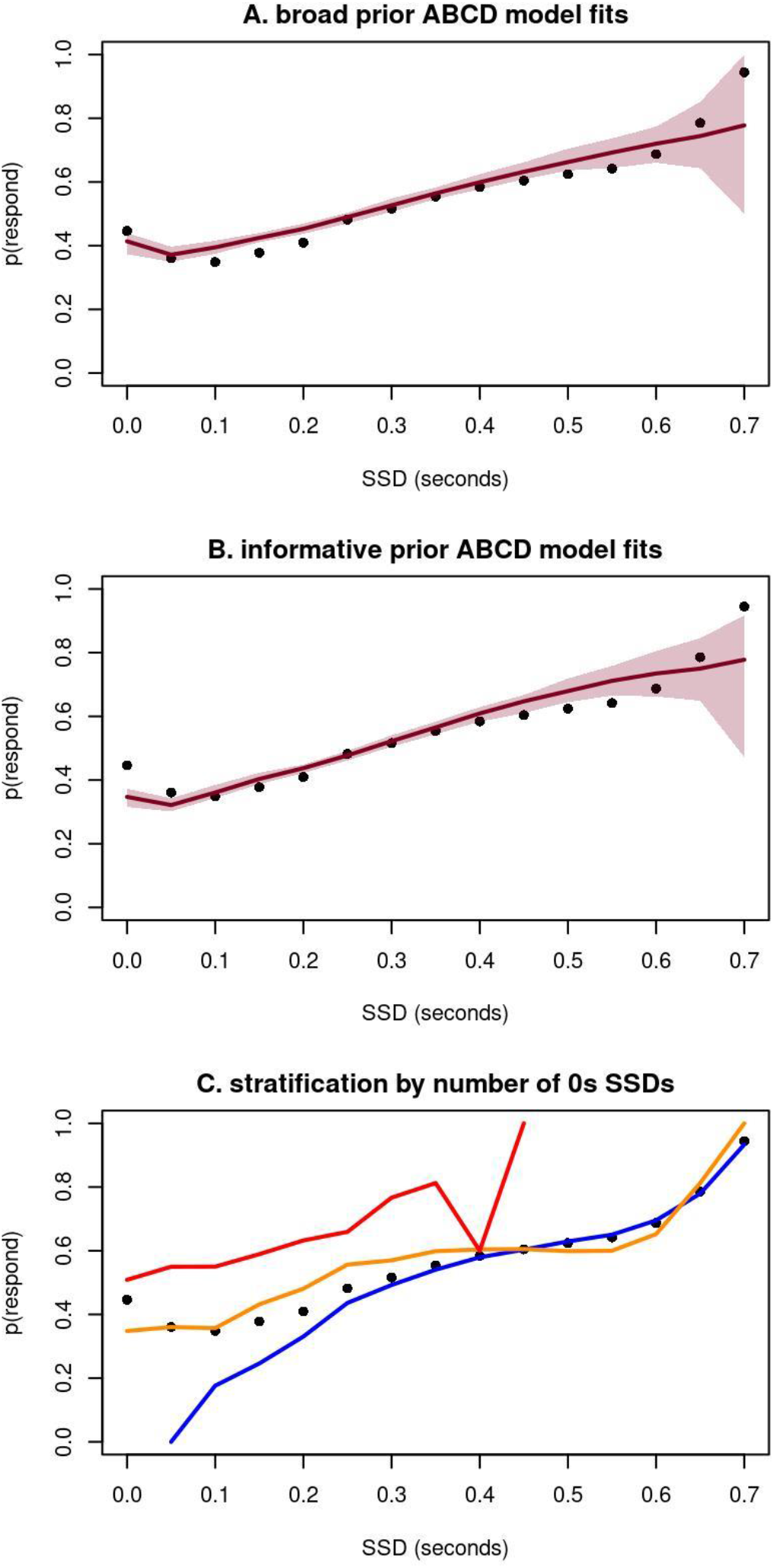
Plots characterizing the inhibition function when trials are binned across participants by specific SSDs. **A**) Posterior predictive plot of the median predictions (red line) and 99% credible interval (CI) of predictions (red shading) of the ABCD stop-signal model when it is fit with broad, uninformative priors, overlayed with empirical data (dots). **B**) Posterior predictive plot of the median (red line) and 99% CI (red shading) of predictions of the ABCD stop-signal model when it is fit with informative priors derived from the hierarchical model fit, overlayed with empirical data (dots). **C**) Plot comparing the group average probability of inhibition at each SSD (dots) with the average probability of inhibition of subgroups stratified by their number of 0s SSD trials: 0 (blue line), 1-9 (orange line) and >=10 (red line)

## Acknowledgements

A.W. was supported by National Institute on Alcohol Abuse & Alcoholism T32 AA007477 (to Frederic Blow) and National Institute on Drug Abuse K23 DA051561. D.M was supported by an Aspasia Budget grant from the Department of Psychology, University of Amsterdam. C.T. was supported by an Advanced ERC grant 743086 UNIFY (to Eric-Jan Wagenmakers). A.H. was supported by Australian Research Council grants DP210100313 and DP200100655 and a Révész Visiting Professor Fellowship at the Department of Psychology, University of Amsterdam.

Data used in the preparation of this article were obtained from the Adolescent Brain Cognitive Development^SM^ (ABCD) Study (https://abcdstudy.org), held in the NIMH Data Archive (NDA). This is a multisite, longitudinal study designed to recruit more than 10,000 children age 9-10 and follow them over 10 years into early adulthood. The ABCD Study® is supported by the National Institutes of Health and additional federal partners under award numbers U01DA041048, U01DA050989, U01DA051016, U01DA041022, U01DA051018, U01DA051037, U01DA050987, U01DA041174, U01DA041106, U01DA041117, U01DA041028, U01DA041134, U01DA050988, U01DA051039, U01DA041156, U01DA041025, U01DA041120, U01DA051038, U01DA041148, U01DA041093, U01DA041089, U24DA041123, U24DA041147. A full list of supporters is available at https://abcdstudy.org/federal-partners.html. A listing of participating sites and a complete listing of the study investigators can be found at https://abcdstudy.org/consortium_members/. ABCD consortium investigators designed and implemented the study and/or provided data but did not necessarily participate in analysis or writing of this report. This manuscript reflects the views of the authors and may not reflect the opinions or views of the NIH or ABCD consortium investigators. The ABCD data repository grows and changes over time. The ABCD data used in this report came from ABCD Release 1.1, DOI 10.15154/1412097.

## Availability of Data and Materials

Data from the ABCD Study are available through the NIH Data Archive (NDA): nda.nih.gov/abcd. Code for all analyses featured in this study is openly available on the Open Science Framework (OSF): osf.io/2h8a7/.

## Footnotes

1 Given the ex-Gaussian can take on negative values we did, however, impose a fixed lower bound of 0.05s on the stop finishing-time distribution to avoid any possibility of implausible values: assuming bounds from 0-0.1s did not have any noticeable effect on our results.

2 We note that when Bissett et al. (2021) used each specific SSD to form “absolute” bins they found that there was an apparent increase in responding on 0s SSD trials relative to other short SSD trials, which is unexpected given race model assumptions. However, we show in Supplemental Materials that this uptick in responding is an artifact of averaging across individuals who have different levels of inhibitory performance. We also show that the ABCD model can account for the uptick when it is unconstrained by priors, likely because this lack of constraint allows the model to be overfit to a subset of individuals with uncommonly poor performance.

3 It should also be noted that the large sample size here detects minor deviations that are less likely to be apparent in more typical experimental studies (e.g., even in relatively large sub-samples of 100 participants, none of the small misfits produced by the BEESTS-ABCD model in Figure 2 are evident).

4 We also point out that trigger failures are already known to bias non-parametric SSRT (Matzke et al., 2017) and, as the BEESTS-ABCD model includes a trigger failure parameter, this parameter can partially account for differences between BEESTS-ABCD and non-parametric SSRT estimates. However, simulation studies reported in this section hold this parameter constant; it is held at a single value, across conditions in which other parameters vary, for each individual featured in Figure 4 and held at a single value across all simulated individuals featured in Figure 5.

5 We also conducted more intensive parameter recovery studies with a subset of individuals’ parameter values that were intended to identify possible biases or second modes in parameter ranges (see Materials and Methods; results available at osf.io/2h8a7/), and again found little evidence for bias.

6 Bissett et al. (2021) also point out potential limitations related to fMRI measurement, namely that the difference in stimulus presentation time between go and stop trials confounds fMRI contrasts that compare these trial types to isolate activity related to inhibition. However, as there are multiple existing confounds already inherent in these conventional stop-signal task contrasts (e.g., stop trials have shorter RTs, additional stimuli presented, and a lower rate of occurrence relative to go trials), we view the ABCD design feature as a minor addition to the more general problems with using subtraction logic in fMRI contrasts, which have been extensively discussed elsewhere (Poldrack & Yarkoni, 2016). Therefore, we think the limitations that the ABCD task design imposes on fMRI measurement are not substantially greater than limitations of fMRI measurement common to all stop-signal tasks.

